# Exofection as a Therapeutic Modality: Restoring P-gp Activity via Trophoblast-Derived EV in Neuroinflammatory Disorders

**DOI:** 10.64898/2026.04.02.716001

**Authors:** Ananth K Kammala, Madhuri Tatiparthy, Sravan Gopalkrishna Shetty Sreenivasmurthy, Klarissa H Garza, Shaneilahi Budhwani, Lauren Richardson, Ramkumar Menon, Balaji Krishnan

## Abstract

**Background:** P-glycoprotein (P-gp/ABCB1) is a key efflux transporter that maintains barrier integrity by clearing xenobiotics and toxic metabolites. At the feto-maternal interface, trophoblast-derived extracellular vesicles (CTC-EVs) naturally and transiently transfer functional P-gp to maternal decidual cells, restoring lost and or reduced P-gp function (exofection) to sustain pregnancy homeostasis. A similar loss of P-gp at the blood brain barrier (BBB) contributes to impaired amyloid-β (Aβ) clearance and neuroinflammation in Alzheimer’s disease. We investigated whether CTC-EV-mediated exofection could restore P-gp function in human brain endothelial cells (hBECs) and enhance Aβ clearance under inflammatory and neurodegenerative conditions.

**Methods:** CTC-EVs were isolated and characterized by nanoparticle tracking analysis and western blotting for P-gp and EV markers. Transcriptomic profiling of CTC-EVs identified enrichment of transporter-related genes, including solute carriers and ABC transporters, along with inflammatory mediators. Network analysis revealed coordinated modules linking EV cargo to transporter regulation, endocytosis/trafficking pathways, and inflammatory remodeling processes converging on BBB efflux activity. hBECs were exposed to LPS (500 ng/mL, 48 h) with or without CTC-EVs. P-gp expression was assessed by immunofluorescence (mean fluorescence intensity, MFI) and western blotting, while functional efflux was measured using Calcein-AM assays. Aβ oligomer transport was evaluated using a transwell hBEC model. In vivo, 3xTg-AD mice received intravenous CTC-EVs (1×10L/day for 5 days), followed by assessment of P-gp expression, Aβ burden, and neuroinflammatory markers. Pharmacokinetic studies in P-gp knockout mice were conducted to confirm functional transporter recovery.

**Results:** LPS exposure significantly reduced P-gp expression in hBECs (41.3% decrease in MFI, p=0.0084), which was restored by CTC-EVs (46.7% increase vs. LPS, p=0.0121). Exofection increased P-gp by a 2.1-fold following EV treatment as determined by western blot. Functional assays demonstrated enhanced efflux, with a 38.5% reduction in intracellular Calcein fluorescence (p<0.001). Network-informed mechanisms supported coordinated regulation of transporter and trafficking pathways. CTC-EVs improved Aβ transport across inflamed hBEC monolayers. In vivo, EV-treated 3xTg-AD mice exhibited increased P-gp expression in the frontal cortex (38.6%) and hippocampus (42.1%), reduced Aβ plaque burden (27.9%), and decreased inflammatory markers (IL-1β and TNF-α, p<0.05). In P-gp knockout mice, EVs reduced brain drug accumulation by 22.4% (p=0.032), confirming restoration of transporter function.

**Conclusion:** CTC derived EVs are natural carriers of functional transporter proteins and restore efflux capacity in compromised endothelial barriers. Integration of transcriptomic and network analyses highlights coordinated regulation of transporter, trafficking, and inflammatory pathways underlying exofection. This reproductive biology inspired strategy offers a promising therapeutic approach for enhancing Aβ clearance and mitigating neuroinflammation in Alzheimer’s disease.

## Introduction

Alzheimer’s disease and related dementias (AD/ADRD) are increasingly significant global health challenges, leading to high levels of disability and dependence among older adults. In the United States, over 7 million people currently live with Alzheimer’s Disease (AD), with annual healthcare and caregiving costs expected to grow from approximately $384 billion in 2025 to nearly $1 trillion by 2050^1^. Despite extensive research over the years, effective disease-modifying treatments remain scarce, a critical unmet need driven by the complex pathophysiology of AD. A major contributing factor is that the key disease processes start years before symptoms appear, and many potential therapies fail to reach or maintain effective sustained levels in the brain^2^, often due to limitations in crossing the blood-brain barrier (BBB) or maintaining therapeutic concentrations within the central nervous system.

A central pathological feature of AD is the accumulation of amyloid-β (Aβ) peptides that aggregate into plaques and trigger downstream synaptic dysfunction, neuroinflammation, and neuronal loss^3^. Although Aβ is produced throughout life as a by-product of amyloid precursor protein processing, its steady-state level in the brain is tightly controlled by clearance pathways rather than production alone^4^. This delicate balance underscores the critical role of efficient Aβ removal. A substantial fraction of Aβ is eliminated across the BBB via coordinated transport mechanisms within the neurovascular unit, including receptor-mediated transcytosis (e.g., LRP1) and ATP-binding cassette transporters such as P-glycoprotein (P-gp/ABCB1) located predominantly on the luminal side of brain endothelial cells^5–7^. Consistent with a clearance-driven model, reduced BBB transporter function and vascular inflammation are increasingly recognized as early contributors to Aβ retention and disease progression^8–10^, highlighting BBB dysfunction as a key therapeutic target.

Multiple, partly convergent mechanisms have been proposed to explain BBB efflux failure in AD. These include (i) impaired LRP1–P-gp functional coupling, (ii) inflammatory signaling (e.g., NF-κB) that suppresses transporter activity, and (iii) enhanced P-gp turnover that occurs through ubiquitin–proteasome-dependent degradation^11–13^. Crosstalk among endothelial cells, astrocytes, and pericytes further modulate the barrier phenotype. Under chronic inflammatory conditions, this intercellular signaling can shift the BBB toward a less protective, more permeable, and less efficient clearance state^14^. An automated image-analysis study of postmortem brain tissue from the National Institute of Health (NIH) and the Karolinska Institute found a 53% reduction in capillary P-gp protein levels in AD patients compared with age-matched controls (p < 0.01), consistent with a role for impaired efflux in human disease states ^15,16^. Pharmacological approaches aiming to induce P-gp expression or activity have shown limited success, frequently constrained by systemic toxicity, inadequate delivery to brain endothelium, off-target drug–drug interactions, and the difficulty of sustaining transporter restoration over time^16^. These significant challenges underscore a fundamental limitation that current strategies often attempt to upregulate endogenous, potentially compromised, protein machinery. These limitations highlight the need for alternate strategies that directly replace or replenish dysfunctional BBB proteins, rather than relying solely on transcriptional upregulation.

Our pregnancy studies investigated how fetal and maternal tissues coordinate physiology to sustain a healthy gestation^17–20^. This work led us to identify an extracellular vesicle (EV)-mediated, cross-barrier communication mechanism between the fetus and the mother^17,21–25^. At the feto-maternal interface, fetal trophoblast cells release EVs that deliver functional molecular cargo to maternal tissues to maintain homeostasis. We term this EV-mediated transfer “exofection”^26,27^. Notably, chorion trophoblast cell-derived extracellular vesicles (CTC-EVs) are uniquely suited for therapeutic applications due to their physiological role in mediating feto–maternal communication, where they naturally transfer bioactive cargo including functional transporter proteins across biological barriers while maintaining immune tolerance and achieving high systemic bioavailability during pregnancy. Importantly, our prior work demonstrating heterogeneity in inflammatory responses at the feto maternal interface revealed that CTCs exhibit a markedly attenuated response to endotoxin compared to maternal decidual cells, with selective activation of stress-response pathways and minimal induction of proinflammatory signaling. This intrinsic immunological resilience positions CTCs as a stable and protected cellular source, capable of generating EVs with preserved functional integrity even under inflammatory conditions. Together, these features highlight CTC-EVs as evolutionarily optimized nanocarriers that combine barrier-crossing capability, low immunogenicity, and sustained functional cargo delivery, making them highly promising for restoring transporter function in compromised systems such as the blood–brain barrier and for developing next-generation therapies targeting inflammation-driven diseases^28^. Their inherent capacity for cross-barrier transport and cargo delivery makes them an ideal candidate for addressing BBB dysfunction in AD. Based on these observations, we hypothesized that trophoblast EVs could deliver functional P-gp to BBB endothelial cells, restore efflux capacity, enhance Aβ handling, and attenuate neuroinflammatory responses associated with AD pathology. To test this hypothesis, we combined mechanistic *in vitro* studies in inflamed human brain microvascular endothelial cells with *in vivo* evaluation in the late-onset mouse model of AD/ADRD (3xTg-AD) at early stages where Aβ precedes tau tangle formation^29,30^ The 3xTg-AD model is particularly valuable for this study as it recapitulates key pathological hallmarks of AD, including amyloid plaque formation and tau pathology, in a progressive manner. By focusing on early stages where Aβ pathology is prominent and precedes tau tangle formation, we can specifically investigate the impact of BBB dysfunction on Aβ clearance and the therapeutic potential of P-gp restoration before widespread neurodegeneration. We assessed EV-mediated restoration of P-gp expression and function, changes in Aβ uptake/clearance-related readouts, and BBB transporter activity using tacrolimus as a P-gp substrate-based functional probe. Together, these experiments introduce exofection as a distinct therapeutic framework for neurodegenerative disease, prioritizing direct, EV-enabled protein replacement at the BBB to restore protective transport and support brain homeostasis.

## Results

Our investigation aimed to characterize circulating tumor cell-derived extracellular vesicles (CTC-EVs) and evaluate their therapeutic potential, particularly concerning P-glycoprotein (P-gp) modulation, in both in vitro models of inflammation and an in vivo model of Alzheimer’s disease (AD).

### Characterization of CTC-Derived Extracellular Vesicles

To establish the fundamental properties of the therapeutic agent, CTC-derived EVs were first rigorously characterized (Figure 1). Nanoparticle tracking analysis (NTA) revealed a predominant vesicle size of 136.8 ± 69.9 nm (Figure 1A), consistent with the typical size range for exosomes. Further confirmation of their exosomal identity was obtained through ExoView analysis, which demonstrated the surface expression of canonical tetraspanin markers CD63, CD81, and CD9 (Figure 1B). Western blot analysis independently validated the presence of these key exosome markers at the protein level (Figure 1C). Quantitative ELISA revealed a dose-dependent enrichment of P-gp within these isolated EVs, with P-gp concentration peaking at 1E+09 particles/mL (Figure 1D). These findings confirm that the isolated CTC-EVs are bona fide exosomes and carry significant levels of P-gp. Thus, these results establish that the identity of the isolated vesicles as exosomes and, importantly, identifies P-gp as a significant cargo, setting the stage for functional investigations.

**Figure 1.**
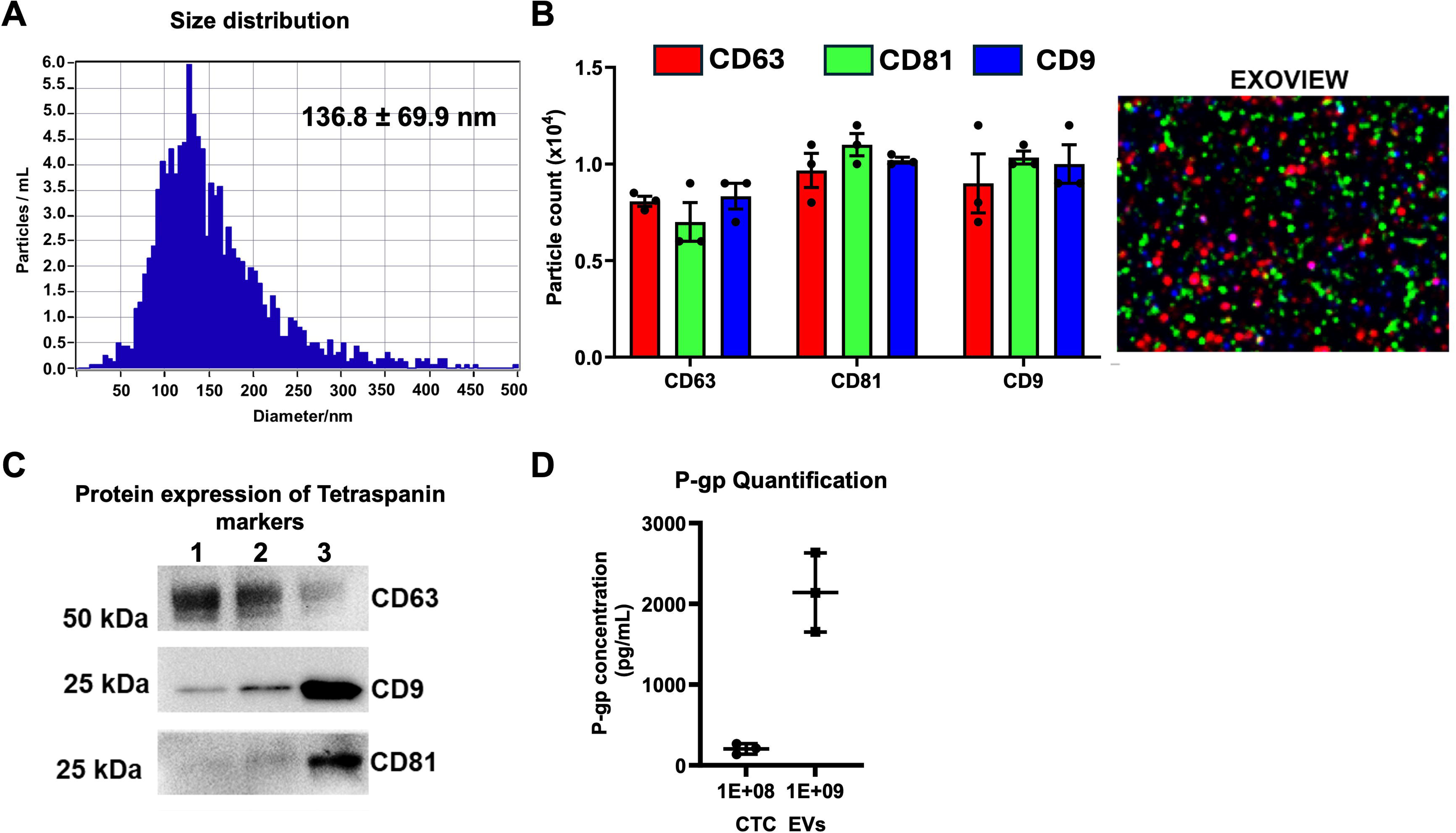
Characterization of CTC-derived extracellular vesicles (EVs). (A) Nanoparticle tracking analysis (NTA) showing a predominant vesicle size around 100 ± 25 nm, consistent with the typical size range for exosomes. (B) ExoView analysis confirming the surface expression of canonical tetraspanin markers CD81, CD63, and CD9, further validating the exosomal nature of the vesicles. (C) Western blot analysis validating the presence and protein expression of exosome markers CD63, CD81, and CD9. (D) ELISA quantification revealing a dose-dependent enrichment of P-glycoprotein (P-gp) in the isolated EVs, with levels peaking at a concentration of 10^9^ particles/mL.

### CTC-EVs Restore P-gp Expression and Efflux Activity in Inflamed Brain Endothelial Cells

Given the presence of P-gp in CTC-EVs, we next investigated their ability to modulate P-gp expression and function in an in vitro model of inflammation, utilizing human brain endothelial cells (hBECs) treated with lipopolysaccharide (LPS) to mimic inflammatory insults (Figure 2 and Figure 3).

**Figure 2.**
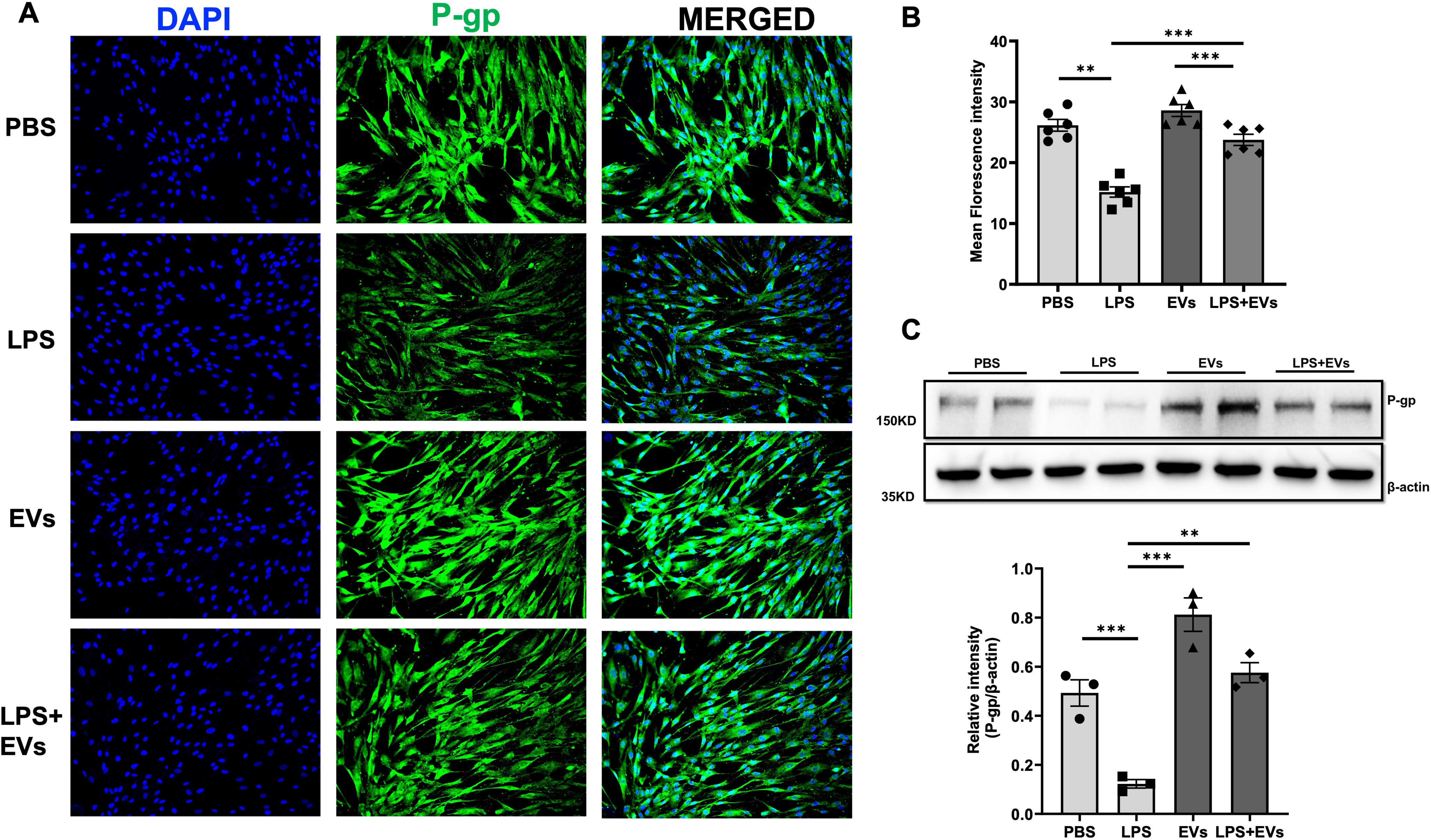
CTC-EVs restore P-gp expression in inflamed brain endothelial cells. Human brain endothelial cells (hBECs) were treated with lipopolysaccharide (LPS; 500 ng/mL, 48 h) to induce inflammation, with or without co-treatment using CTC-derived extracellular vesicles (CTC-EVs; 1x10 particles/mL). (A) Immunocytochemistry images showing reduced P-gp (green) expression after LPS treatment, which is notably preserved in cells co-treated with LPS and CTC-EVs; nuclei are counterstained with DAPI (blue). (B) Quantification of P-gp fluorescence intensity, confirming the LPS-induced downregulation of P-gp and its partial recovery upon co-treatment with CTC-EVs. (C) Representative Western blots illustrating P-gp protein levels across different treatment groups. (D) Densitometry analysis of Western blots, further supporting the CTC-EV-mediated rescue of P-gp expression, with P-gp signal normalized to β-actin as a loading control.

**Figure 3.**
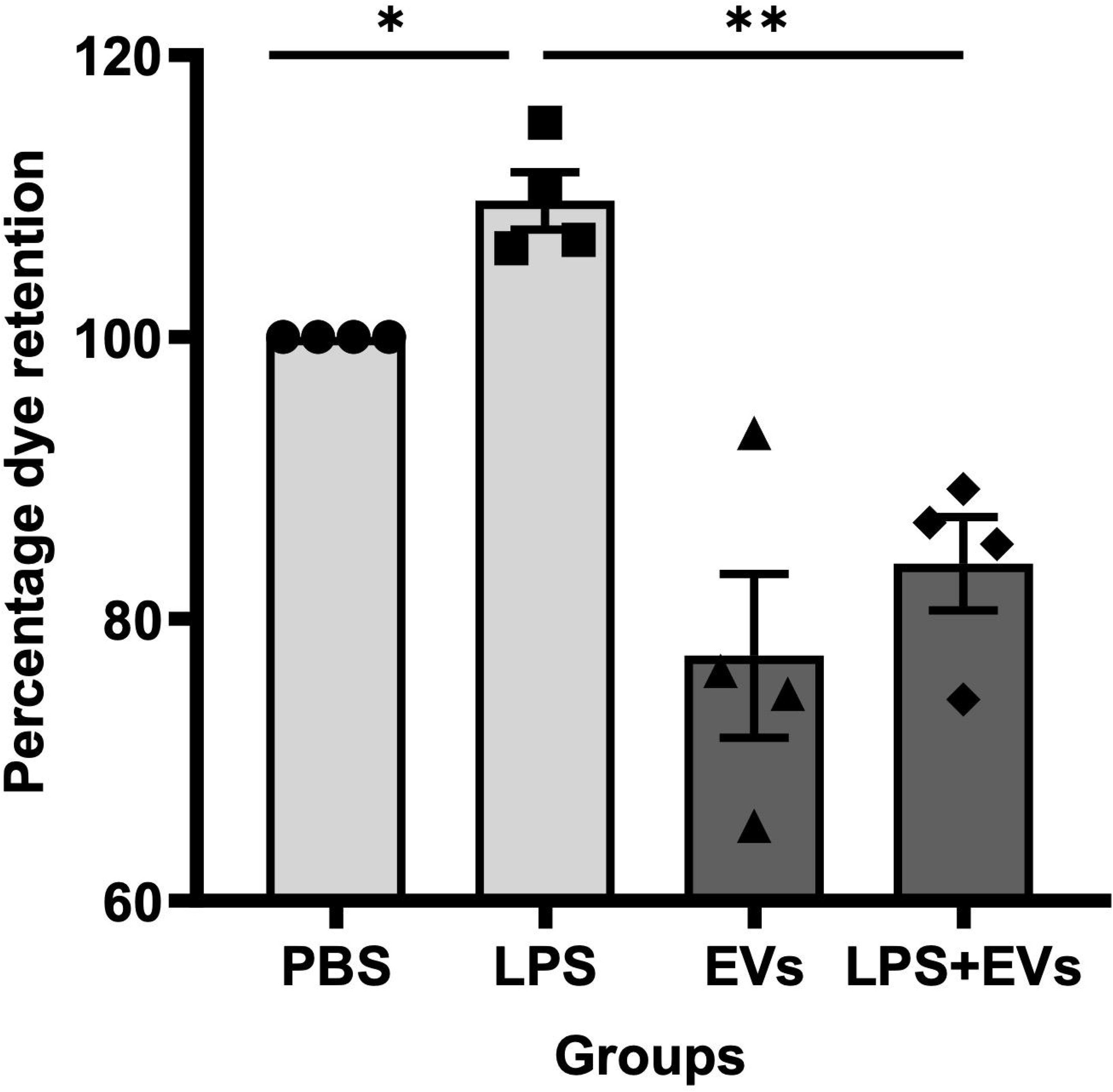
Restoration of P-gp efflux activity in inflamed hBECs by CTC-derived EVs. (A) Schematic illustration of the P-gp efflux assay. This assay demonstrates that cells with active P-gp efficiently extrude a hydrophobic fluorescent dye, resulting in reduced intracellular fluorescence. Conversely, cells with less or inactive P-gp retain the dye, leading to higher intracellular fluorescence. Fluorescence measured at 520 nm is inversely proportional to P-gp activity. (B) Quantitative results of the P-gp efflux assay. LPS treatment significantly reduced efflux function (*p < 0.05 vs. PBS), indicating impaired P-gp activity. Co-treatment with CTC-EVs significantly increased efflux compared to LPS alone (**p < 0.01), confirming a functional recovery of P-gp activity.

LPS treatment (500 ng/mL for 48 hours) significantly reduced P-gp expression in hBECs, as evidenced by immunocytochemistry (Figure 2A) and quantified by a significant decrease in mean fluorescence intensity (Figure 2B). Importantly, co-treatment with CTC-EVs (1x10 particles/mL) preserved P-gp expression, with levels appearing similar to untreated controls and significantly higher than LPS-only treated cells (Figure 2A, 2B). These findings were further corroborated by western blot analysis, which showed a marked reduction in P-gp protein levels following LPS treatment, a reduction that was significantly rescued by co-treatment with CTC-EVs (Figure 2C, 2D). Beyond expression, we assessed the functional activity of P-gp using a fluorescence-based efflux assay (Figure 3). The assay principle relies on the inverse relationship between intracellular fluorescent dye retention and P-gp efflux activity. Consistent with the reduced expression, LPS treatment significantly impaired P-gp efflux function, leading to a higher percentage of dye retention compared to untreated cells (*p < 0.05 vs. PBS). Remarkably, co-treatment with CTC-EVs significantly restored P-gp efflux activity, resulting in a lower percentage of dye retention compared to LPS-treated cells (**p < 0.01), indicating a functional recovery of the P-gp pump (Figure 3). Thus, our results collectively demonstrate that CTC-EVs effectively counteract LPS-induced inflammation by restoring both the expression and, critically, the functional efflux activity of P-gp in brain endothelial cells.

### CTC-EVs Restore Aβ Oligomer Transport Across Inflamed Brain Endothelial Cells

Building on the restoration of P-gp function, we investigated whether CTC-EVs could mitigate the impact of a neuropathological agent, specifically amyloid-beta (Aβ) oligomers, in an in vitro model of the blood-brain barrier (BBB) (Figure 4). Human brain endothelial cells (hBECs) were cultured on Transwell inserts and exposed to LPS to induce inflammation, followed by the addition of Aβ oligomers (AβO) to the apical chamber (Figure 4A). ELISA quantification of AβO levels in the basal compartment revealed that LPS treatment significantly reduced AβO translocation across the endothelial layer compared to control conditions (*p < 0.05). This suggests that inflammation impairs the BBB’s ability to transport AβO.The addition of CTC-EVs rescued this effect, significantly enhancing AβO transport under inflammatory conditions (Figure 4B). This indicates that CTC-EVs can restore the compromised AβO clearance mechanism in an inflamed BBB model. Thus, our studies provide direct evidence that CTC-EVs can restore the transport of toxic Aβ oligomers across inflamed brain endothelial cells, highlighting a potential mechanism for mitigating AD pathology.

**Figure 4.**
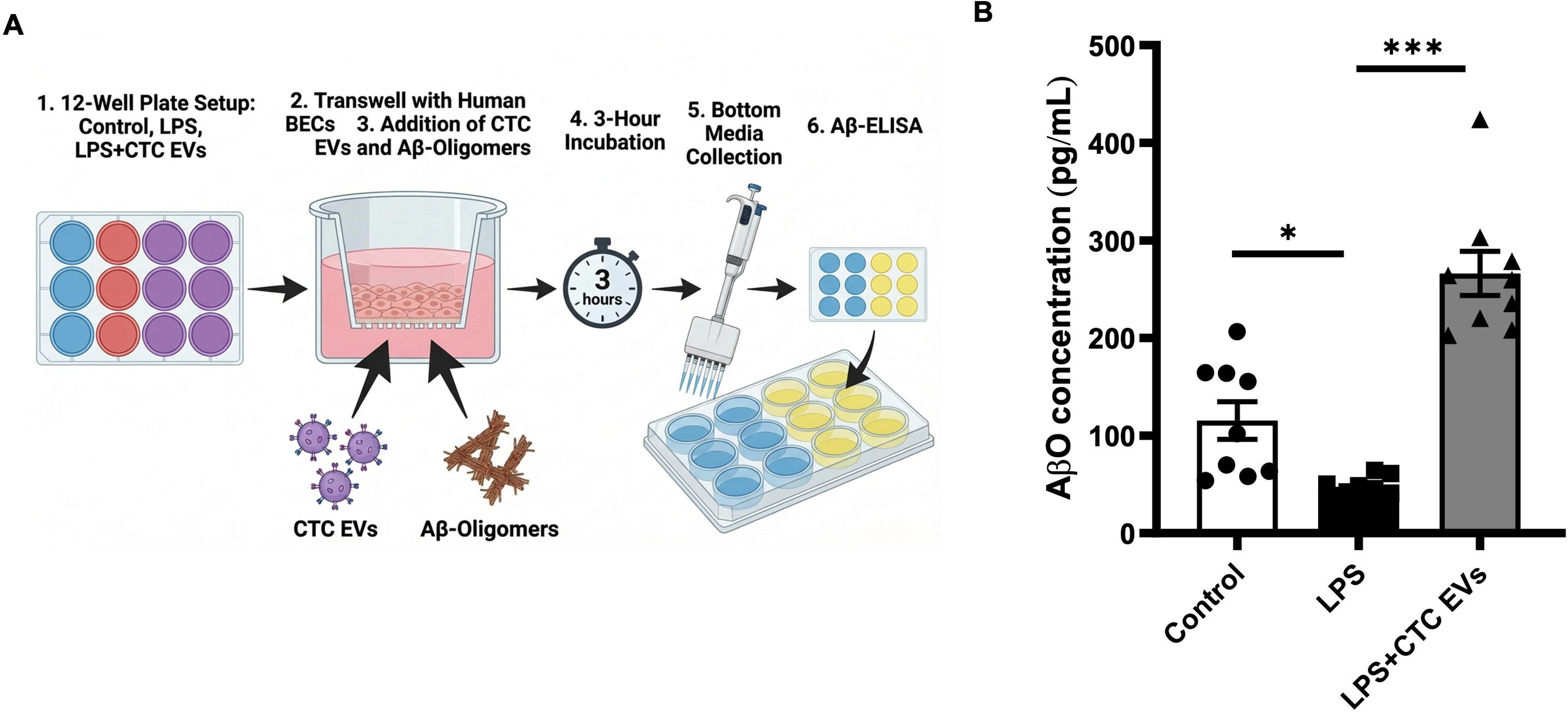
CTC-derived EVs restore Aβ oligomer transport across inflamed brain endothelial cells. (A) Schematic of the experimental workflow. Human brain endothelial cells (BECs) were cultured on Transwell inserts and subjected to various treatments: Control, LPS, or LPS + CTC-EVs. Amyloid-beta (Aβ) oligomers were subsequently added to the apical chamber, and the basal media were collected after 3 hours to quantify Aβ transfer using ELISA. (B) ELISA quantification of Aβ oligomer (AβO) levels in the basal compartment. LPS treatment significantly reduced AβO translocation across the endothelial layer compared with the control group. Importantly, the addition of CTC-EVs rescued this effect and enhanced AβO transport under inflammatory conditions. Data are presented as mean ± SEM; *p < 0.05.

### CTC-EV Transcriptomics Reveals Integrated Transporter and Inflammatory Networks

Transcriptomic profiling of CTC-EVs followed by network analysis identified a complex interaction landscape (Supplementary Fig. S1) encompassing genes involved in transcriptional regulation, stress responses, extracellular matrix organization, and inflammation. Prominent clusters included inflammatory mediators (e.g., IL1β, TNF, ICAM1) and structural/cytoskeletal genes, indicating diverse functional cargo.

A focused network (Figure 5) highlighted three major functional modules relevant to barrier function: (i) a transporter module comprising ABC transporters (ABCB1, ABCC family, ABCG2) and solute carriers (SLC family), (ii) a trafficking/endocytosis module involving regulators such as LRP2, PIK3CA, and GNA12, and (iii) an inflammatory remodeling module including C3, PLAU, IL32, ICAM1, and AXL. These modules converged on BBB efflux activity, suggesting coordinated regulation of transporter function. Notably, transporter-associated genes were closely linked to trafficking pathways, supporting a mechanism in which EV cargo facilitates both delivery and functional integration of efflux machinery. The concurrent presence of inflammatory mediators indicates that CTC-EVs may also modulate the inflammatory microenvironment.

**Figure 5.**
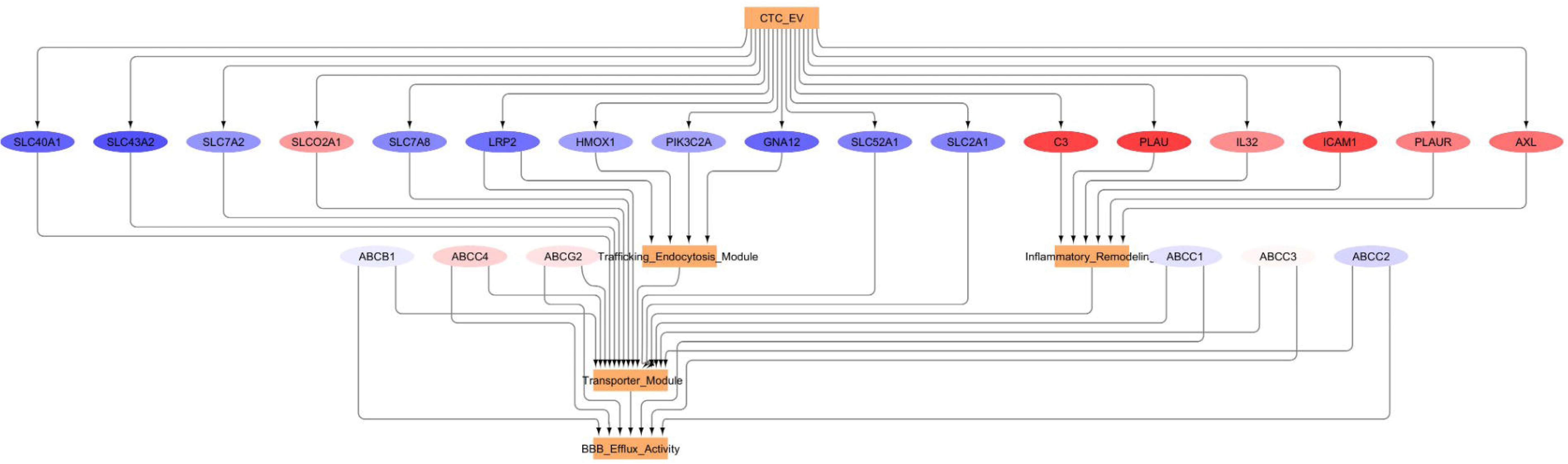
Focused network of CTC-EV transcriptomic cargo. Network analysis of CTC-EV–associated genes highlighting key functional modules. Nodes represent genes (red: upregulated; blue: downregulated), and edges indicate interactions. Major modules include transporter (ABC and SLC families), trafficking/endocytosis (e.g., LRP2, PIK3CA), and inflammatory signaling (e.g., C3, ICAM1). These converge on BBB efflux activity.

### P-gp Deficit in 3xTg-AD Mice and In Vivo Restoration by CTC-EVs

To translate our in vitro findings to a more complex biological system, we utilized the 12-month-old 3xTg-AD mouse model, which progressively accumulates Aβ and tau pathologies, leading to synaptic dysfunction and memory deficits.

First, we established the baseline P-gp profile in this model (Supplementary Figure 2). Western blot analysis of brain homogenates from 12-month-old 3xTg-AD mice revealed a significant decrease in P-gp levels in both the frontal cortex and hippocampus compared to age-matched wild-type controls. This finding confirms a critical P-gp deficit in key brain regions of the AD model, providing a strong rationale for therapeutic intervention.

Subsequently, we investigated the in vivo efficacy of CTC-EVs in modulating P-gp expression in the 3xTg-AD mouse model (Figure 6). Twelve-month-old 3xTg-AD mice received acute intravenous administration of CTC-EVs, and P-gp expression was assessed in the frontal cortex and hippocampus via Western blotting (Figure 6A, 6B). Quantification of P-gp expression, normalized to total protein, showed that while 3xTg-AD mice exhibited reduced P-gp levels in both regions compared to C57BL/6J controls, acute CTC-EV treatment led to a significant improvement in the P-gp profile specifically in the frontal cortex (p < 0.01, n = 3-6 per group) (Figure 6C). Notably, no significant improvement in P-gp levels was observed in the hippocampus following this acute treatment regimen. After establishing that there is a (Supplementary Figure 1) a critical P-gp deficit in the 3xTg-AD mouse model, we were able to provide evidence for the therapeutic potential of our findings (Figure 6) where acute administration of CTC-EVs partially restores P-gp expression in a region-specific manner within the brain of 3xTg-AD mice, with a significant effect observed in the frontal cortex.

**Figure 6.**
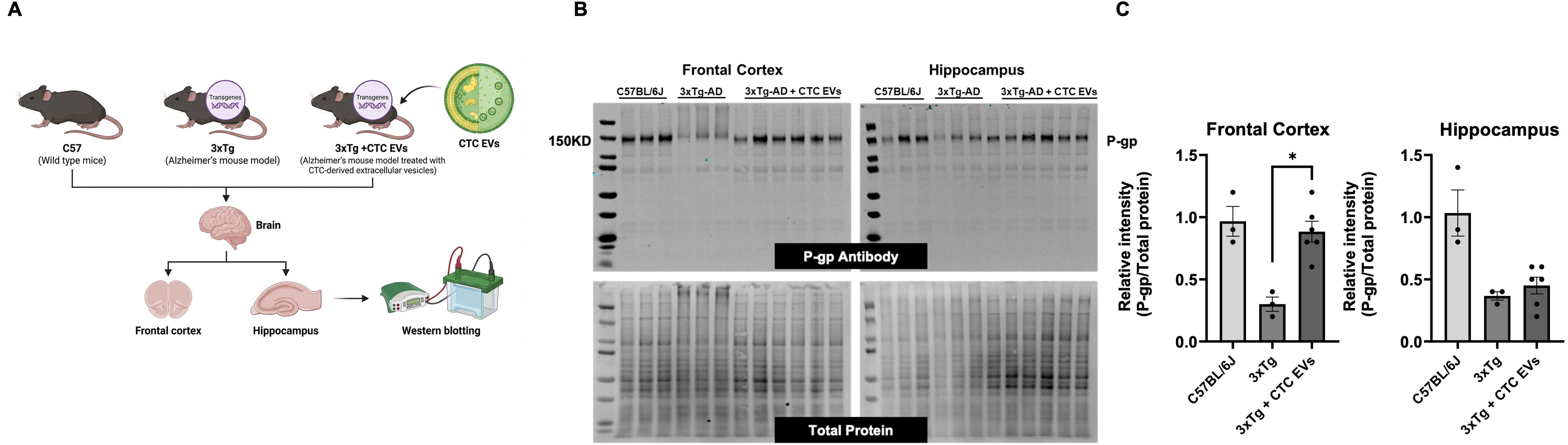
CTC-derived EVs restore P-glycoprotein (P-gp) expression in Alzheimer’s 3xTg-AD mouse model. (A) Schematic representation of the in vivo experimental design. Twelve-month-old 3xTg-AD mice were acutely administered CTC-derived extracellular vesicles (CTC-EVs) intravenously. Following treatment, brains were collected for regional analysis, with the frontal cortex and hippocampus dissected, homogenized, and processed for Western blotting to assess P-gp expression. (B) Representative Western blots illustrating P-gp protein levels in whole-brain homogenates from the frontal cortex and hippocampus. Groups include wild-type C57BL/6 mice, untreated 3xTg-AD mice, and 3xTg-AD mice treated with CTC-EVs. Total protein staining served as the loading control. (C) Quantification of P-gp expression normalized to total protein. 3xTg-AD mice exhibited a marked reduction in P-gp levels in both the frontal cortex and hippocampus compared with wild-type controls. Treatment with CTC-EVs partially restored P-gp expression, with a significant increase observed specifically in the frontal cortex (p < 0.01, n = 3-6 per group).

### CTC-EVs Restore Functional P-gp–Mediated Drug Efflux in P-gp Knockout Mice

To evaluate the impact of EV-mediated transporter delivery on tissue-level drug disposition, we utilized the P-gp knockout (KO) mouse model and assessed Tacrolimus concentrations following CTC-EV administration (Figure 7A). Our previous work established that CTC-derived EVs restore functional P-gp activity in vivo through exofection. Building on this, the current analysis focuses specifically on tissue distribution patterns.

**Figure. 7:**
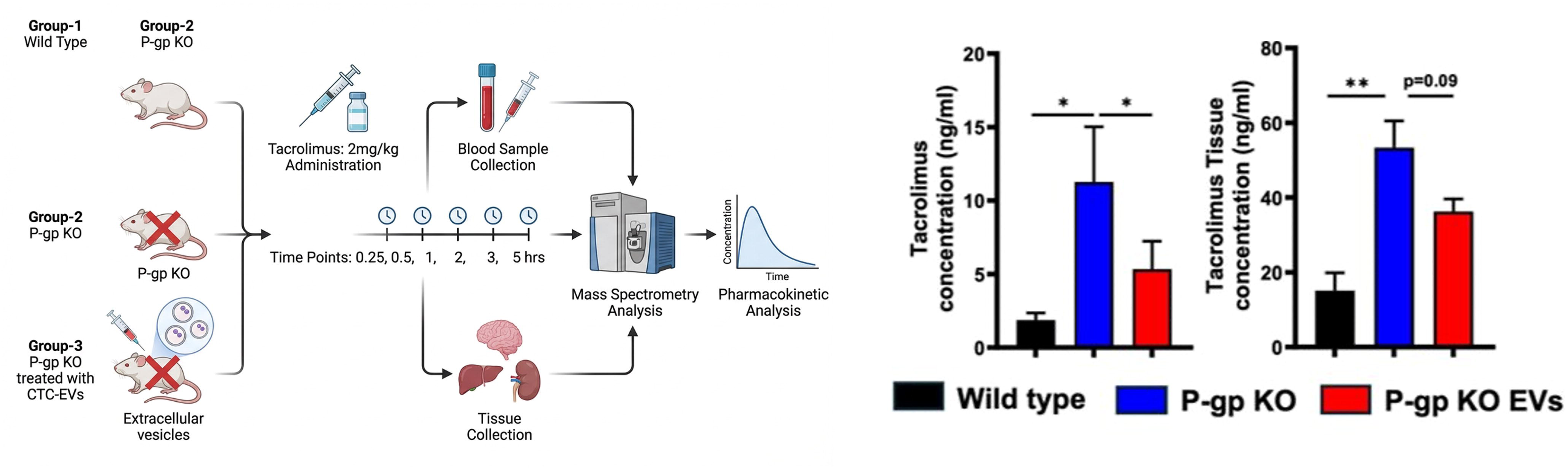
Restoration of P-gp Function in P-gp KO Mice by CTC-EVs. (A) Schematic representation of experimental design to determine the functional effect of CTC-EVs in P-gp KO mice by evaluating pharmacokinetics of Tacrolimus. (B) Tacrolimus plasma concentration over time in wild-type (WT), P-gp KO, and KO+EV-treated mice, showing reduced drug clearance in KO mice and partial restoration of P-gp function in KO+EV mice. (C) Tissue distribution of Tacrolimus across the Brain and uterus demonstrated higher drug accumulation in KO mice and reduced levels in KO+EV mice, indicating restored efflux activity. (D) Pharmacokinetic parameters comparing WT, KO, and KO+EV groups support that CTC-EVs enhance P-gp mediated drug transport. Data were analyzed using an unpaired t-test, and values are presented as mean + SEM (N = 3 per group).

As expected, KO mice exhibited elevated Tacrolimus accumulation in P-gp–sensitive organs, particularly the brain and uterus, reflecting impaired efflux capacity (Figure 7B). In contrast, KO mice treated with CTC-EVs showed a marked reduction in Tacrolimus concentrations within these tissues compared to untreated KO controls. This decrease in tissue drug burden suggests that EV-delivered P-gp contributes to localized restoration of efflux activity at barrier sites.

These findings complement our prior demonstration of exofection-mediated transporter rescue by showing that CTC-EVs can also influence organ-level drug distribution. The observed reduction in Tacrolimus accumulation in the absence of endogenous P-gp supports the concept that EV-derived transporter cargo is functionally integrated and capable of modulating tissue-specific efflux dynamics. These results collectively establish the therapeutic potential of CTC-derived EVs for conditions characterized by P-gp deficiency or dysfunction.

## Discussion

This study identifies a previously unrecognized therapeutic mechanism in which CTC EVs restore BBB efflux capacity through the direct delivery (exofection) of fully functional P-gp. By colllopting an evolutionarily refined fetal–maternal communication system, these findings show that vesicle-based transporter transfer can reconstitute BBB function and signaling networks that are typically refractory to pharmacologic rescue. The work broadens transporter biology by demonstrating that P-gp activity in recipient endothelial cells can be re-established through direct, vesicle-mediated protein delivery rather than relying on transcriptional activation or de novo gene expression.

Inflammation-induced repression of P-gp at the BBB emerges from these data as a critical node where neuroimmune signaling converges on barrier dysfunction^14^. LPS-driven innate immune activation, used here to model chronic neuroinflammatory states relevant to AD, markedly reduces endothelial P-gp abundance and efflux capacity in a manner consistent with TLR4–NF κB-mediated destabilization of ATP-binding cassette (ABC) transporters, alterations in membrane lipid organization, and enhanced proteasomal degradation^31–33^. Clinical and experimental studies have reported loss or mislocalization of P-gp in cerebral microvessels in AD, which is associated with impaired Aβ clearance and progressive accumulation of Aβ peptide in the parenchyma, supporting a mechanistic link between reduced P-gp function and disease progression^34^. Pharmacologic approaches using anti-inflammatory agents, such as ibuprofen^35^, as well as metabolic modulation of endothelial cells, have previously suggested that augmenting P-gp activity can remodel BBB transport properties in neurodegenerative settings. Here, CTC-EVs act as molecular supplements that circumvent transcriptional silencing by delivering fully assembled, membrane-embedded P-gp directly to brain endothelial cells^36,37^. The vesicle-derived transporter incorporates into recipient plasma membranes in a conformation competent for ATP hydrolysis, substrate binding, and active efflux, thereby restoring P-gp function under conditions in which small-molecule inducers or conventional gene-delivery strategies have shown limited ability to re-establish BBB transporter activity.^38–41^

The rescue of Aβ translocation across inflamed endothelial monolayers further shows that EV-mediated P-gp replacement restores an important aspect of BBB function^42,43^. P-gp is a key efflux mechanism for monomeric and oligomeric Aβ species; therefore, inflammatory suppression of P-gp is likely to speed up parenchymal Aβ buildup, promote plaque formation, and increase microglial-driven neurotoxicity^44,45^. Our findings that CTC-EVs normalize Aβ transport even with LPS indicate that these exosome-transporters remain responsive to substrate flux and can participate in physiological Aβ clearance pathways that are otherwise impaired early in AD development. This supports a model in which fetal trophoblast EV biology offers a natural way to protect tissues from toxic peptide buildup, a mechanism that can be repurposed for neurodegeneration.

Importantly, transcriptomic and network analyses of CTC EV cargo provide a systems-level framework supporting this mechanism. The EV transcriptome is enriched in transporter-associated genes, trafficking regulators, and inflammatory mediators, which organize into interconnected modules linking ABC transporters and solute carriers with endocytic pathways and inflammatory signaling networks. These data suggest that exofection is not solely a protein transfer event but part of a coordinated program that facilitates transporter delivery, membrane integration, and functional stabilization. The close association between transporter genes and trafficking pathways supports a model in which EV cargo enhances both the availability and intracellular routing of efflux machinery, while concurrent inflammatory mediators may modulate the local microenvironment to favor restoration of barrier function.

The in vivo restoration of P-gp in the frontal cortex of 3xTg-AD mice strengthens the translational potential of this approach, highlighting that vesicle-mediated delivery can overcome the chronic transporter deficits that occur during AD progression. The frontal cortex, one of the first regions to show breakdown of the blood-brain barrier and inflammatory stress^46^, demonstrated the strongest recovery, indicating regional differences in EV uptake, membrane turnover rates, or susceptibility to transport restoration. Restoring P-gp in these brain areas suggests a capacity to re-establish efflux function in circuits where impaired clearance of neurotoxic substances speeds up cognitive decline^43^. These results support the emerging view that repairing the BBB is essential for effective AD treatment and that restoring efflux may complement Aβ-targeted immunotherapies by reducing parenchymal load and lowering ARIA risk.

Pharmacokinetic experiments in P-gp deficient mice primarily served as a physiological validation that CTC-EVs can deliver functional transporters to target tissues. In this stringent background, EV treatment partially restored P-gp dependent handling of a probe substrate, indicating that vesicle-delivered P-gp can become functionally active in vivo and support the feasibility of using CTC-EVs to correct transporter deficits at the BBB.

Taken together, these findings support a mechanistic model in which CTC-EVs function as endogenous, precisely targeted vesicles that have evolved for barrier reinforcement^28^. Their tetraspanin-rich membrane structure enables fusion with endothelial cells; the enclosed P-gp arrives fully folded and glycosylated; and the EV surface provides chaperoning lipids that facilitate transporter insertion into recipient membranes^47,48^. This efficient delivery system avoids ER-dependent folding bottlenecks and bypasses inflammatory transcriptional repression, establishing exofection as a fundamentally distinct mode of protein delivery compared to gene therapy or nanoparticle methods. The fetal origin of these vesicles is especially notable: the feto-maternal interface requires efficient transfer of protective transporters during inflammatory stress, similar to challenges observed at the BBB in neurodegeneration. AD pathology may therefore be mitigated by adopting strategies originally evolved to maintain pregnancy-specific barrier homeostasis.

Important limitations warrant cautious interpretation. The observed partial restoration of P-gp in the hippocampus, in contrast to the stronger recovery in the frontal cortex, highlights potential regional constraints on transporter incorporation or stability. This disparity may stem from differences in EV uptake efficiency, varying membrane turnover rates across distinct brain regions, or the potentially greater susceptibility of the hippocampus to severe or chronic neuroinflammation, which could impede the therapeutic effect. These regional variations, alongside the need for systematic definition of CTC-EV biodistribution, the durability of transporter expression, and the immunological consequences of repeated administration, underscore critical areas for future research. Furthermore, the complex cargo of EVs necessitates careful evaluation for potential off-target delivery of signaling molecules to non-vascular tissues, and successful clinical translation will ultimately require scalable GMP-compliant manufacturing with standardized potency assays.

Despite these challenges, this work outlines a mechanistically grounded platform for BBB repair. By showing that transporter deficits can be reversed via endogenous vesicle-based exofection, the study positions CTC-EVs as a tractable modality to correct BBB failure across AD, Parkinson’s disease, vascular dementia, and chemotherapy-related cognitive impairment. This also motivates future efforts to engineer EVs carrying combinatorial cargo such as anti-inflammatory microRNAs, multi-transporter panels, or tight-junction stabilizers. More broadly, the data suggest that regenerative principles operating at the feto-maternal interface can be harnessed to restore neurovascular homeostasis, providing a rationale for clinical translation of EV-mediated transporter replacement in neurodegenerative disease.

## Conclusion

This study identifies trophoblast-derived EV as a potent, naturally occurring platform for restoring blood–brain barrier efflux function through exofection of fully active P-glycoprotein. By demonstrating that EV-delivered P-gp (i) integrates into endothelial membranes, (ii) reestablishes ATP-dependent transport, and (iii) normalizes clearance of both xenobiotics and Aβ species under inflammatory stress, we establish a mechanistic framework in which transporter deficiency in neurodegenerative disease can be reversed without genetic modification. The ability of these vesicles to rescue efflux capacity in AD/ADRD models and its verification in P-gp null animals underscore the robustness of this mechanism and highlight a previously unrecognized mode of BBB repair. These findings reveal that principles governing fetal–maternal barrier protection can be strategically repurposed to combat neurovascular dysfunction, offering a new class of biologically informed therapeutics for AD and other disorders characterized by impaired transporter signaling. By restoring a key regulatory node that coordinates neuroimmune activation, toxic peptide clearance, and CNS pharmacokinetics, EV-mediated exofection presents a transformative approach to enhance brain resilience and reshape treatment strategies across neurodegenerative diseases.

## Materials and methods

### Cell culture

Fetal membrane CTCs were obtained from term individuals, not in labor (TNIL), and immortalized using the HPV16 E6E7 retrovirus method^49,50^. CTCs were cultured in a specialized DMEM/Ham’s F12 medium containing penicillin, streptomycin, bovine serum albumin, heat-inactivated fetal bovine serum (HI-FBS), and ITS-X, along with CHIR99021, A83-01, SB431542, L-ascorbic acid, valproic acid (VPA), Y27632, and epidermal growth factor to support their proliferation and maintenance.

Human brain endothelial cells (hBECs; ATCC, Manassas, VA, USA) were cultured in endothelial growth medium (EGM-2, Lonza) supplemented with 10% heat-inactivated fetal bovine serum (FBS; Gibco), 1% penicillin–streptomycin (Gibco), and growth factors according to the manufacturer’s instructions. Cells were maintained at 37 °C in a humidified incubator with 5% CO₂ and used between passages 4 and 8. For all experiments, cells were seeded to reach confluence and allowed to stabilize for 24 hours before treatment^49,50^.

### EV isolation by the tangential flow filtration (TFF) system

Our team optimized the isolation and purification of EVs using the TFF protocol^51^. Culture media from CTCs at 70-80% confluency were collected and stored at −80°C. To pre-clean the media before EV isolation, sequential centrifugation was performed: 300 *g* for 5 minutes to remove cell contamination, 3000 *g* for 10 minutes to eliminate cell debris, and 17,000 *g* for 15 minutes to discard microvesicles, followed by filtration through a 0.2 µm membrane to retain only particles smaller than 0.2 µm. The TFF system was then prepared with an exosome membrane module. The system was first flushed with water through both the retentate and filtrate tubes, sterilized with 0.5M NaOH, and rinsed with water. The pre-cleaned media was introduced into the sample chamber, where flow rate (30-40 mL/min), pressure (20-30 psi), and room temperature were carefully maintained to ensure stable filtration. The TFF 100 K membrane filtrate effectively removed particles smaller than 100 kDa, including free soluble proteins. Finally, the purified EV sample was collected and further concentrated into a small volume using Beckman ultracentrifugation.

### Exosome size distribution and concentration measurement by Zetaview

Exosomes concentration and size distribution were analyzed with the ZetaView™ PMX 110 (Particle Metrix, Meerbusch, Germany) with 8.02.28 software. The Zetaview machine was prepared by washing the main line and sample injection line with water (3X) before use. The instrument calibration was done with the beads at a 1:250000 dilution. Frozen Exosome samples were gently thawed on ice and then diluted in filtered water at a 1:1000 ratio. This dilution allowed for accurate measurement of particle concentration (particles/mL) and average size distribution for each sample. To ensure the instrument’s accuracy, it was thoroughly rinsed with filtered distilled water in between each sample analysis.

### P-gp quantification by ELISA

P-gp levels in exosome preparations were quantified using a Human Plllgp/ABCB1 ELISA kit (Reddot, Cat. RDRlllPgplllHu) according to the manufacturer’s instructions, with minor modifications. Briefly, the kit reagents were equilibrated to room temperature, and 100 µL of standards or exosome samples (normalized by protein content, as determined by the BCA assay) were added to antibody-coated wells and incubated overnight at 4 °C. The following day, plates were washed three times with 350 µL wash buffer, incubated with 100 µL biotinylated anti–Plllgp detection antibody for 60 min at 37 °C, washed five times, and then incubated with 100 µL HRPlllstreptavidin conjugate for an additional 60 min at 37 °C. After five further washes, 90 µL TMB substrate was added and the reaction was allowed to develop for 10–20 min at 37 °C in the dark before addition of 50 µL stop solution, and absorbance was read immediately at 450 nm using a microplate reader; sample concentrations were interpolated from the standard curve.

### P-gp and tetraspanin marker characterization by western blot analysis

For characterization of Plllgp and tetraspanin markers, CTClllEVs were lysed in radioimmunoprecipitation assay (RIPA) buffer containing 50 mM TrislllHCl (pH 8.0), 1% Triton Xlll100, 1 mM EDTA (pH 8.0), 150 mM NaCl, and 0.1% SDS supplemented with phenylmethylsulfonyl fluoride and protease/phosphatase inhibitor cocktails. Lysates were cleared by centrifugation at 12,000 g for 20 min at 4 °C, and protein concentration in the supernatant was determined using the Pierce BCA assay (Thermo Scientific). Equal amounts of protein were separated on 10% SDS–polyacrylamide gels and transferred to PVDF membranes using a BiolllRad wet transfer system. Membranes were blocked for 2 h at room temperature in 5% nonfat dry milk in TBS with 0.1% Tweenlll20 (TBST) and incubated overnight at 4 °C with primary antibodies against the EV markers CD63, CD81, and CD9 (1:1000, System Biosciences, EXOABlllkitlll1) and Plllgp (1:1000, Abcam, ab170904). After four washes with TBST, membranes were incubated for 1 h at room temperature with HRPlllconjugated secondary antibodies (antilllrabbit for Plllgp, 1:10,000, Amersham NA934VS; secondary antibodies for CD markers supplied with EXOABlllkitlll1, 1:5000), washed again, and developed using an enhanced chemiluminescence detection system (ECL Max, Amersham).

### Treatment of hBECs with LPS and CTC EVs

hBECs were seeded at a density of 0.20 × 10 cells per well in six-well plates and cultured for 70–80% confluency. For the inflammatory model, cells were treated with 500 ng/mL LPS (LPS; Sigma, Cat# L2880), prepared from a 1 mg/mL stock solution. Cells were exposed to LPS for 48 hours to induce inflammation. For exofection experiments, cells were co-treated with 500 ng/mL LPS treatment and exosomes at a concentration of 1 × 10¹ particles for 48 hours.

### Immunocytochemistry

After the LPS treatment, the cells were first fixed and permeabilized with 4% PFA and 0.5% Triton X and blocked with 3% BSA in PBS. The cells were then incubated overnight at 4°C with a primary antibody targeting P-gp (Abcam, Cat# ab235954), diluted at 1:300. Following thorough PBS washes, the cells were incubated for 1 hour with a secondary antibody, Alexa Fluor 488-conjugated anti-rabbit IgG (Abcam, Cat# ab150073), at a dilution of 1:1000. After additional PBS washes, the cells were stained with NucBlue™ Fixed Cell Ready Probes™ reagent (Invitrogen) and then mounted using Mowiol (Calbiochem Cat# 475904) mounting medium.

hBECs were grown on collagen-coated glass coverslips, treated as above, washed with PBS, and fixed with 4% paraformaldehyde for 15 min at room temperature. Cells were permeabilized with 0.1% Triton X-100 in PBS for 10 min, blocked in 5% bovine serum albumin (BSA) for 1 h, and incubated overnight at 4 °C with primary antibody against P-glycoprotein (P-gp; clone C219 or equivalent, 1:100; Cell Signaling Technology or Abcam). After washing, samples were incubated with Alexa Fluor–conjugated secondary antibodies (Invitrogen, 1:500) for 1 h at room temperature and counterstained with DAPI (1 µg/mL) before mounting with antifade medium. Images were acquired using a laser-scanning confocal microscope (e.g., Zeiss LSM 880) with identical acquisition settings across conditions, and mean fluorescence intensity of P-gp per field was quantified using ImageJ/Fiji.

### Western blotting

For Western blotting, treated hBECs were rinsed with ice-cold PBS and lysed in RIPA buffer (Thermo Fisher) supplemented with protease and phosphatase inhibitors (Roche). Lysates were cleared by centrifugation at 14,000 g for 15 min at 4 °C, and protein concentration was determined using the BCA assay (Pierce). Equal amounts of protein (20–30 µg) were resolved on 7.5–10% SDS–polyacrylamide gels and transferred to PVDF membranes (Millipore). Membranes were blocked in 5% nonfat dry milk in TBS-T (Tris-buffered saline with 0.1% Tween-20) for 1 h, incubated overnight at 4 °C with antibodies against P-gp (1:1,000) and β-actin (1:5,000; Sigma-Aldrich) as a loading control, then probed with HRP-conjugated secondary antibodies (1:5,000; Jackson ImmunoResearch) for 1 h at room temperature. Bands were visualized by enhanced chemiluminescence (ECL; GE Healthcare) and imaged on a digital imager (e.g., Bio-Rad ChemiDoc). Densitometry was performed using ImageJ, and P-gp signal was normalized to β-actin; data are presented as relative intensity values in the violin plots

### P-gp efflux assay

Plllgp efflux activity was assessed using the Multidrug Resistance Assay Kit (Cayman Chemical, 600370) following the manufacturer’s instructions. Briefly, treated cells were washed with PBS and incubated at 37 °C for 30 min in 100 µL culture medium containing vehicle or Verapamil (Plllgp inhibitor, 1:2000). Cells were then exposed to 100 µL calceinlllAM working solution (2 µL stock in 10 mL medium) for an additional 30 min at 37 °C, after which the supernatant was removed by centrifugation at 400 g for 5 min and cells were washed twice with cold medium. Fresh cold medium (200 µL) was added, and intracellular calcein fluorescence was measured at 485 nm excitation and 535 nm emission using a microplate reader; higher fluorescence corresponds to reduced Plllgp efflux activity.

### Transwell assay

Human brain microvascular endothelial cells (HBMECs; ATCC) were seeded onto collagenlllcoated 0.4lllµm Transwell inserts (24lllwell, Corning) and grown to confluence in EGMlll2 medium with 10% FBS and 1% penicillin–streptomycin at 37 °C in 5% CO₂, then exposed for 24 h to either vehicle (control), 500 ng/mL LPS, or LPS plus CTClllderived extracellular vesicles (CTClllEVs). After equilibration in HBSS, synthetic human Aβ oligomers were added to the apical chamber and, following a 3 h incubation, basal medium was collected and Aβ levels were quantified by ELISA according to the manufacturer’s instructions, with data expressed as Aβ concentration (pg/mL).

### CTC-EV Transcriptomics and Network Analysis

RNA was isolated from purified CTC-derived extracellular vesicles using a low-input RNA extraction kit and quality assessed by Bioanalyzer. Libraries were prepared using a stranded total RNA-seq kit and sequenced on an Illumina platform. Reads were quality filtered, aligned to the human genome (GRCh38), and gene counts generated. Differential expression analysis was performed using DESeq2, with adjusted p-value <0.05 considered significant.

Differentially expressed genes were analyzed using STRING for protein–protein interactions and visualized in Cytoscape. Functional modules and hub genes were identified based on network topology. Pathway enrichment (GO/KEGG) was performed using clusterProfiler (FDR <0.05). A focused network highlighting transporter, trafficking, and inflammatory pathways was derived from enriched gene sets.

### Animal studies

Animal procedures were followed in accordance with the Institutional Animal Care and Use Committee (IACUC) at the University of Texas Medical Branch, Galveston, under approved protocol number 041107F.

Twelvelllmonthlllold male 3xTglllAD mice and agelllmatched C57BL/6 wildllltype controls (Jackson Laboratory) were housed under standard conditions with ad libitum access to food and water, and all procedures were approved by the institutional animal care and use committee. 3xTglllAD mice received CTClllderived extracellular vesicles (CTClllEVs) by intravenous injection at a dose of 1E+09 particles once daily for 2 consecutive days; wildllltype and untreated 3xTglllAD mice served as controls. Twentylllfour hours after the final injection, mice were deeply anesthetized, perfused with cold PBS, and brains were rapidly removed on ice; frontal cortex and hippocampus were dissected, snaplllfrozen in liquid nitrogen, and stored at −80 °C until protein extraction. Band intensities were quantified using ImageJ, Plllgp signal was normalized to total protein for each lane, and data from frontal cortex and hippocampus were expressed as relative Plllgp levels for C57BL/6, 3xTglllAD, and 3xTglllAD + CTClllEV groups (n = 3 per group).

### P-gp functional activity in the P-gp KO mice

Timed-pregnant FVB/NTac wildllltype mice (Taconic Biosciences) and Plllgp knockout mice (FVB.129P2lllAbcb1a Abcb1b N12) were housed in a temperaturelllcontrolled facility, and pregnancy was confirmed by detection of a vaginal sperm plug and subsequent weight gain. Dams were randomly assigned to three groups: wild type (WT), Plllgp knockout (KO), and Plllgp knockout treated with CTClllderived EVs (KO + CTClllEVs). On embryonic day 14 (E14), mice in the KO + CTClllEV group received two intravenous doses of CTClllEVs (1 × 10¹ particles per dose) administered 24 h apart, whereas WT and KO groups received vehicle. On E16, all three groups were given tacrolimus (2 mg/kg, intraperitoneal), and maternal blood and tissues were collected at 15, 30, 60, 120, 180, and 300 min to assess pharmacokinetics, with a particular focus on brain tacrolimus levels. At each time point, animals were euthanized, and brain, blood, lungs, and reproductive tissues were harvested for analysis, allowing comparison of tacrolimus disposition between WT, KO, and KO + CTClllEV mice and evaluation of CTClllEV–mediated restoration of Plllgp–dependent efflux at the BBB.

### Mouse tissue processing

After the experiment, the isolated tissue was frozen. Then, the tissue was thawed, and the samples were weighed on the glass slide. They were then kept at −80°C for 30 minutes. Then, the tissue was finely cut, and PBS was added in a 1:4 ratio. Finally, half a spoonful of 1mm Zrsio beads was added. Using the bullet blender, the tissue was homogenized until no tissue remained. The sample was collected and stored at −80°C.

### Analytical method for Tacrolimus measurement

A liquid chromatography-tandem mass spectrometry (LC-MS/MS) method with electrospray ionization (ESI) was developed and validated for quantifying tacrolimus in mouse plasma, placenta, fetal membrane, uterus, and lung samples following FDA guidelines. Tacrolimus-*d3* was selected as the internal standard. The chromatographic separation of tacrolimus and its internal standard was performed on a Waters XTerra® MS C18 HPLC column (50 × 2.1 mm, 3.5 µm) at 45°C. The mobile phase consisted of acetonitrile + water with 2% NH₄OH (v/v), and gradient elution was used at a flow rate of 350 µL/min. An API 4000 triple quadrupole mass spectrometer, equipped with a Turbo V ion source, operated in negative mode. Quantification was conducted using multiple reaction monitoring (MRM) with transitions of m/z 802.5 → 560.6 for tacrolimus and m/z 805.5 → 563.6 for tacrolimus-*d3*. The calibration range for tacrolimus in mouse serum ranged from 0.26 ng/mL to 44.8 ng/mL. Method accuracy ranged from 92% to 112%, with relative standard deviation (RSD) values not exceeding 11%. Quality control samples at high, medium, and low concentrations were analyzed alongside test samples to ensure result reliability.

### Statistical analysis

Data were analyzed as mean values with the standard error of the mean (SEM) indicated. Two-tailed Student’s t-tests were utilized to compare the two groups. When the analysis involved more than two groups, one-way ANOVA was performed, employing a general linear model with a univariate analysis approach. More than three groups were analyzed using two-way ANOVA with multiple comparisons. All the experiments were performed with three biological replicates or three different experiments.

## Author contributions

Conceptualization: AK

Methodology: AK, BK

Investigation: AK, MRT, SM, KG, SB

Visualization: LR, RM, BK

Funding acquisition: AK

Project administration: AK, RM, BK

Supervision: AK, LR, RM, BK

Writing – original draft: AK, BK

Writing – review & editing: SM, BK, LR, RM

## Data and materials availability

All data are available in the main text or the supplementary materials.

## Figure Legends

**Supplementary Figure 1. Global interaction network of CTC-EV transcriptome**

Comprehensive network of differentially expressed genes in CTC-EVs. Nodes represent genes (red/blue indicate expression changes), and edges denote interactions. The network shows broad functional diversity, including inflammatory, structural, metabolic, and regulatory pathways.

**Supplementary Figure 2. Baseline P-gp levels in 3xTg-AD mouse brain regions.**

This figure demonstrates that P-glycoprotein (P-gp) levels are significantly decreased in both the frontal cortex and hippocampal homogenates from 12-month-old 3xTg-AD mice compared to age-matched control animals. This establishes a key pathological deficit in P-gp expression within these critical brain regions in the Alzheimer’s disease model, providing a rationale for therapeutic intervention.

