## Supplementary figures and images for "Exofection as a Therapeutic Modality: Restoring P-gp Activity via Trophoblast-Derived EV in Neuroinflammatory Disorders"

### Supplementary figure 1

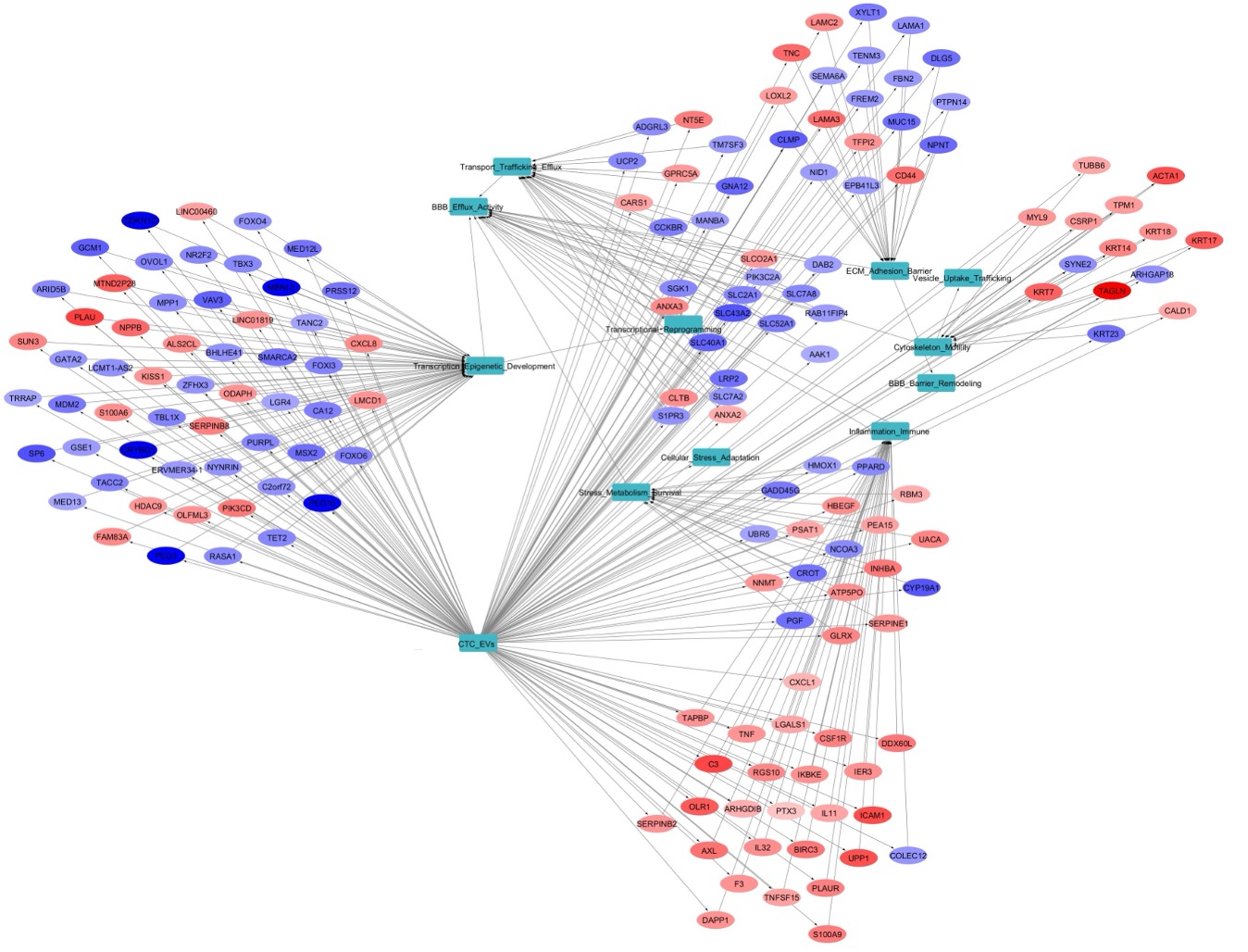

### Supplementary figure 2

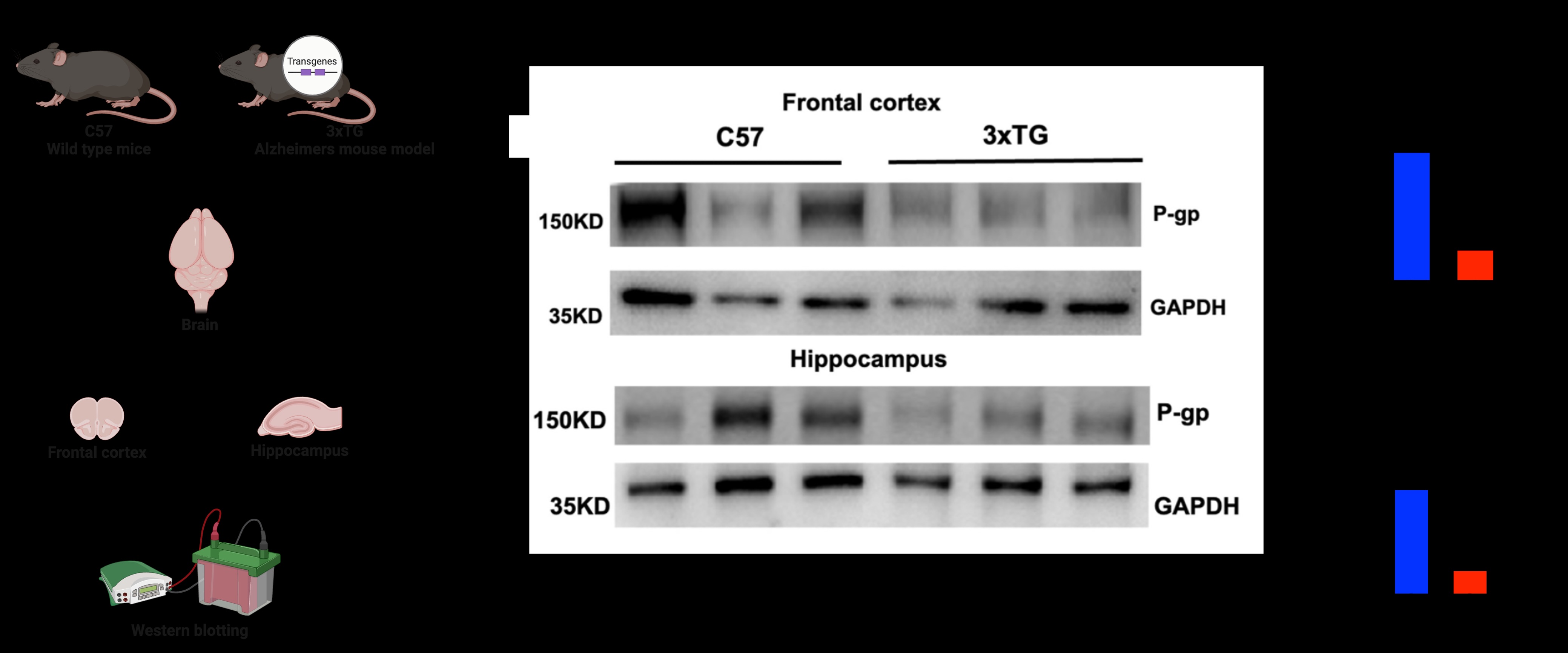
